# Immunomodulatory effects of Simvastatin in Toll-like receptor-2 stimulated Rheumatoid arthritis synoviocytes

**DOI:** 10.1101/2022.05.30.493863

**Authors:** Mohammad Saeed, Zeiyad Al-Karakooly, Nawab Ali, Robert A. Ortmann

## Abstract

Fibroblast-like synoviocytes (FLS) are major effector cells in Rheumatoid arthritis (RA). Toll-like receptor-2 (TLR2) activation of FLS plays a key role in RA pathogenesis. Recently, statins have been shown to modulate RA. We investigated the immunomodulatory effects of simvastatin (SMV) in vitro, in FLS cell lines from control and RA subjects, in the presence of TLR2 activation. Normal and RA FLS cultures (1×10^5^ cells/well) were treated with SMV then activated with TLR2 ligand (Pam_3_CSK_4_ 100μg/ml) for 24-hours. SMV caused dose-dependent cytotoxicity in both cell lines as assessed by Alamar-blue assay and increased ROS production measured by flow cytometry. Gene expression of anti-oxidants including GSR, TXNRD1, PRDX1, SOD2 and MT3 was upregulated. BNIP3 expression was increased 2200-fold in RA-FLS after SMV/TLR2 treatment pointing to apoptosis as a cause of observed cytotoxicity. Apoptosis increased in both cell lines with SMV to 25%. Akt decreased dose-dependently in RA FLS assessed by western blots. IP-1, IP-2 and IP3 decreased with SMV/TLR2 treatment indicating cell survival while IL-6 increased dose-dependently. Thus, SMV induces apoptosis in FLS by upregulating BNIP3 and inhibiting Akt. FLS react to survive by increasing IL-6, decreasing IP-1 and IP-2 and upregulating anti-oxidant genes. SMV has immunomodulatory properties and may be used in RA as adjunctive therapy in combination with other more potent disease-modifying medications.

## Introduction

Rheumatoid Arthritis (RA) is a chronic inflammatory disease that is characterized by synovial inflammation and destruction of joint cartilage and bone [1]. Within the joint, RA synovium presents a striking picture of chronic inflammation, with extensive infiltration by macrophages and lymphocytes, and activation of synoviocytes, which are major players in disease pathogenesis [1, 2]. Hyperplasia of the synovial lining in RA is primarily due to proliferation of fibroblast-like synoviocytes (FLS) due to their impaired apoptosis [2]. Moreover, FLS frame a synovial microenvironment that not only produces but also responds to pro-inflammatory cytokines and chemokines, reactive oxygen species (ROS), and matrix metalloproteinases (MMPs) [2, 3]. Together, these cellular and molecular events perpetuate a state of non-resolving inflammation for patients with RA that ultimately result in joint destruction [4]. Several lines of evidence suggest that Toll-like receptors (TLRs) and their endogenous ligands contribute to inflammation, while blockade of TLR-mediated signaling may yield considerable clinical benefit [5 - 7]. Increased expression of TLR-2, 3, 4, and 7 has been reported in RA synovium suggesting an active role for innate immunity in joint inflammation [8-11]. In particular, TLR2 plays a crucial role in RA pathogenesis [7, 8].

Statins are competitive inhibitors of hydroxymethylglutaryl (HMG)-CoA reductase which catalyzes the conversion of HMG-CoA to mevalonate, the rate-limiting step in cholesterol biosynthesis. Statins have a variety of direct anti-inflammatory and immunomodulatory effects not accounted for by their lipid-lowering properties [12, 13]. Mevalonate, through alternate synthetic pathways, also serves as the necessary substrate for the synthesis of several other biologically important lipid intermediates including geranylgeranylpyrophosphate (GGPP) and farnesylpyrophosphate (FPP), involved in the activation of small GTP-binding proteins Rho, Rab and Ras [12]. Statins therefore also have the potential to modulate intracellular signal transduction pathways. It has been shown that statins induce apoptosis in FLS by inhibition of FPP and GGPP synthesis [14], reduce ROS formation by NADPH oxidase inhibition in endothelial cells [15] and decrease secretion of pro-inflammatory cytokines such as IL-6, IL-8, TNF-α and MCP-1 in monocytes [16].

In a murine collagen-induced arthritis (CIA) model, intra-peritoneal treatment with simvastatin (SMV) inhibited the progression of arthritis [13, 17]. Similarly, in a rat CIA model, oral atorvastatin treatment inhibited arthritis and was accompanied by decreased tissue concentrations of various chemokines [18]. Another study, however, was unable to demonstrate any preventive benefit in a murine CIA model using atorvastatin, rosuvastatin or SMV irrespective of the mode of administration [19].

There have been some clinical trials in humans examining the effects of statins on RA [20, 21]. A modest beneficial effect of statins has been demonstrated with decreased RA disease activity [20-24], decreased CRP levels [23] and reduced number of swollen joints [20, 21]. In contrast, a large US medical and pharmacy claims study showed no beneficial effect of statins in more than thirty-one thousand RA patients as measured by initiation and cessation of oral steroids [25]. A similar study from UK also showed lack of clinical benefit [26], while another large population-based cohort study found association with reduced risk of developing RA [27]. Collectively, these studies point to discrepancies in immunomodulatory effects of statins that need further examination, both *in vitro* and *in vivo*.

The objective of the present study is to examine the immunomodulatory properties of SMV in normal and RA-FLS under conditions of TLR-2 activation *in vitro*, as this mimics the environment in the inflamed synovium more closely [7, 8]. Most of the studies aimed at understanding the role of innate immune cells and TLR function in systemic inflammatory diseases such as RA, have largely focused on macrophages, dendritic cells and other cells of myeloid origin. We investigated the immunomodulatory effects of SMV on FLS cytotoxicity and metabolism, ROS production and gene expression, apoptosis, and downstream signaling events that may help to extend our understanding of the pleiotrophic effects of SMV on synoviocytes.

## Materials and Methods

### Reagents

Normal synovial fibroblast cell line (catalog # 4700) was purchased from ScienCell, Carlsbad, CA while the RA synovial fibroblast cell line (catalog # 2646) was from Cell Applications, San Diego, CA. Synoviocyte Medium (ScienCell, #4701-prf), Accutase (Innovative Cell Technologies, San Diego, CA). Fetal calf serum (FCS) and antibiotics were obtained from American Type Culture Collection (ATCC). Trypan blue (Life technologies, Grand Island, NY), Simvastatin (catalog # 10010344; Cayman Chemical, Ann Arbor, MI), TLR2-ligand Pam_3_CSK_4_ (Invivogen, San Diego, CA), Alamar-blue (Life technologies, Grand Island, NY), Annexin V-FITC (Santa Cruz biotechnology, CA), dihyroethidine (DHE; Molecular probes, Eugene, OR, USA), TNF-α and IL-6 ELISA kits (BD Biosciences, San Diego, CA) were obtained. BNIP3 antibody was obtained from Abcam (ab38621,Cambridge, MA), Akt and SOD1 antibodies from Cell Signaling Technology,Danvers, MA and Actin antibody from Sigma-Aldrich, St. Louis, MO. [^3^H]-myo-inositol (specific activity 20 Ci/mmol) was from American Radiolabeled Chemicals and AG-1×8 resin was from BioRad Laboratories. All other biochemicals were obtained from Sigma-Aldrich. All plastic-ware including 75cm^2^ cell culture flasks, 12-well and 96-well plates were purchased from BD Biosciences, San Diego, CA.

### Preparation of Simvastatin (SMV)

SMV 10mg (Cayman Chemical # 10010344) was dissolved in 1ml of DMSO and aliquoted in 50μl. At the time of each experiment one vial was dissolved in 12ml of complete Synoviocyte medium to give a concentration of 100μM (DMSO concentration 0.21%). This working solution was used for treatment of FLS.

### Cell Culture

FLS cell lines (n=2) derived from healthy (ScienCell # 4700) and RA (Cell Applications # 2646) subjects were obtained commercially and were maintained in Synoviocyte Medium (ScienCell) supplemented with 2% fetal calf serum, penicillin / gentamicin (1%) and synovial growth factors. FLS were cultured in 75cm^2^ flasks at 37°C in a humidified chamber under 5% CO_2_ in air. Passage 4 was used for all experiments. Adherent FLS were gently lifted using Accutase (Innovative Cell Technologies). Cell viability (>95%) was assessed using Trypan blue cell-exclusion assay and a hemocytometer was used to count cells to adjust the concentration to 1×10^5^ cells/ml. Unless otherwise stated, cells were grown in 1ml of medium in 12-well plates and pre-treated with SMV (5-50μM) for 4-hours, subsequently washed and then activated by TLR2 ligand (Pam_3_CSK_4_, 100μg/ml) for 24-hours. Basal measurements were provided by medium-only control wells for both cell lines. Treated controls were FLS stimulated with TLR2 ligand only for 24-hours. All experiments were done in triplicates.

### Cell cytotoxicity assay

Alamar-blue, a nontoxic, cell permeable colorimetric redox indicator (Life technologies, Grand Island, NY) was used to measure cell cytotoxicity as previously described [28]. Reducing power of the mitochondrial NADH/NADPH-dependent dehydrogenase in living cells causes Alamar-blue to change from oxidized (blue) form to reduced (pink) form that is directly proportional to cell number. The absorbance of Alamar-blue at 570nm was used as cell viability indicator. Normal and RA FLS (5×10^4^ cells/ well) were seeded into 96-well plates in triplicates overnight. After media change, FLS were treated as described above. Alamar-blue was added during the last 1-hour of incubation period at 37°C and OD^570^ was measured in an ELISA plate reader.

### Flow cytometry analysis of intracellular ROS production

To assess ROS production intracellularly, cells were incubated with 2.5 μM dihyroethidine (DHE; Molecular probes) for 15 min at 37°C as per treatment protocol described above. Cells were subsequently harvested using Accutase, triplicates were pooled and hydroethidine (HE) fluorescence was assessed by flow cytometry. HE fluorescence is an indicator of superoxide radical generation in cells [28]. A minimum of 10,000 events were collected by gating using the side-scatter function for flow cytometric analysis and the results were presented as percentage of gated cells.

### ROS-associated Gene Expression

Total RNA was extracted from FLS using RNeasy kits (Qiagen Inc, Valencia, CA) according to the manufacturer’s protocol. The experiments were carried out in triplicate as described above and the cell lysates were pooled. RNA was quantitated using Nanodrop 2000. cDNA was prepared from 0.5μg RNA used as template with the RT^2^ First Strand kit (Qiagen Inc, Valencia, CA). Gene expression analysis of ROS associated genes (n=84) was performed using the human “oxidative stress and antioxidant defense” PCR Profiler Array (catalog # PAHS-065) from SA Biosciences. Real-time PCR was performed on ABI 7900 HT (Applied Biosystems, Calsbad, CA) according to the manufacturer’s instructions. Thermocycler parameters were 95°C for 10 min, followed by 40 cycles of 95°C for 15 s and 60°C for 1 min.

### Annexin V assay

Apoptosis in FLS cultures was measured using a commercial flow cytometry kit (Santa Cruz biotechnology) for Annexin V-FITC binding to phosphatidylserine [28]. Briefly, cells were harvested from culture plates using Accutase, washed twice in cold PBS followed by addition of 500μl 1X binding buffer (containing optimal concentration of calcium) supplied by the manufacturer. To this, Annexin V-FITC was added per manufacturer’s protocol and cells were incubated for 15-minutes at room temperature before being analyzed by flow cytometry. At least 10,000 events were collected by gating using the side-scatter function in FACS Calibur (Becton Dickinson, San Jose, CA) and the data were analyzed using CellQuest software.

### Cytokine ELISA

Pro-inflammatory cytokines tumor necrosis factor-α (TNF-α) and interleukin-6 (IL-6) were measured in tissue culture supernatants using a commercially available sandwich ELISA kit (BD Biosciences). As recommended by the manufacturer, both negative and positive controls were included in each assay. Standard curves were drawn using cytokine standards as per manufacturer’s recommendations. Experiments were done in triplicates. Supernatants were stored at −70°C and measured at the same time by ELISA to avoid variation in assay conditions.

### Western blots

Cell lysates were prepared by first washing the attached cells once with ice-cold PBS, and then lysed for 15 min with a M-PER Mammalian protein extraction reagent (Pierce Biotechnology, Rockford, IL) containing a protease inhibitor cocktail (Sigma-Aldrich, St. Louis, MO) and 1 mM p-amidinophenyl methanesulfonylfluoride (PMSF). Cell debris was removed by centrifugation at 14,000 g for 5 minutes. Protein concentration in cell lysates was determined by using a standard Coomassie Bradford protein assay (Bio-Rad, Hercules, CA). For SDS-PAGE cell lysates were mixed with an equal volume of 2X SDS sample buffer (50 mMTris, pH 6.8, 4% SDS, 200 mM DTT, 2% glycerol, 0.01% bromophenol blue), and boiled for 2 minutes. Equal amounts of proteins were separated by 10% SDS-PAGE. The proteins from the gels were electrophoretically transferred onto polyvinylidene fluoride (PVDF) membranes (Millipore, Billerica, MA). The membranes were blocked with 5% skim milk prepared in Tris-buffered saline (150 mM NaCl and 20 mM Tris–HCl, pH 7.2) containing 0.05% Tween-20 (TBS-T) for 1-hour at room temperature. The membranes were then incubated with diluted primary antibodies (as per manufacturer’s recommendations) overnight at 4°C with constant shaking. The membranes were washed three times at 5 minutes each with TBS-T and were incubated for 2-hours with a 1:10,000 dilution of an appropriate secondary antibody conjugated to horseradish peroxidase in 2% skim milk in TBS-T (Cayman Chemical Company, Ann Arbor, MI). Signal detection was achieved with a Super Signal West Pico Chemiluminescence kit (Pierce Biotechnology, Rockford, IL) and high performance chemiluminescence film (Amersham Biosciences, Piscataway, NJ).

### Inositol phosphate (IP) measurements

FLS in 12-well plates (1×10^5^ cells/well) were labeled with a 10 μCi [^3^H]-myo-inositol/ml (American Radiolabeled Chemical, Inc., St. Louis, MO) in Synoviocyte medium for 24-hours. Subsequently, the experiments were carried out as described above and the triplicates pooled. The cells were then washed three times with PBS to remove extracellular [^3^H]-myo-inositol. Soluble IPs liberated from cells were extracted as detailed previously [29, 30]. Briefly, the adherent cells were dissolved in 1% SDS to extract total phosphoinositol lipids. For separation into individual IPs (1 to 4), cell debris was removed by centrifugation and clear supernatants were diluted 10-fold and subjected to anion exchange chromatography on gravity fed columns using AG-1 9 8 resin (formate form, 100–200 mesh size, Bio-Rad Laboratories, Hercules, CA). Fractions containing inositol mono, bis, tris, and tetrakis phosphates were separated with increasing concentrations of ammonium formate in a batchwise elution as previously described [31]. Radioactivity was counted in LS6500 Beckman Coulter beta scintillation counter (Beckman Coulter, Inc., Fullerton, CA). Changes in cellular levels of IPs were calculated by normalization with total membrane bound inositol phospholipids from corresponding wells.

### Statistics

All data are presented as means ± SD. Statistical differences between groups were analyzed by using two-tailed unpaired Student *t* test and ANOVA as indicated; *p* < 0.05 was considered statistically significant. For data analysis of qPCR array, SA Biosciences integrated web-based software package for the PCR Array System was used. The software automatically performs all ΔΔC_t_ based fold-change calculations from uploaded raw threshold cycle data obtained from ABI 7900 HT obtained using the ABI Prism software (Applied Biosystems, Calsbad, CA). Briefly, relative changes in gene expression were calculated using the comparative threshold cycle (ΔΔC_t_) method which involves normalizing the C_t_ (threshold cycle number) against average C_t_ of the housekeeping genes on the array, followed by squaring the difference between the normalized average C_t_ of treated and control groups. The result is expressed as fold change.

## Results

### Cell cytotoxicity

To assess cytotoxicity by Alamar-blue, cell concentration standards were prepared by serial dilution of FLS. Cells were plated in duplicate ranging from 5×10^3^ to 2×10^5^ cells /ml in a 12-well plate and cultured for 24-hours in medium. Alamar-blue was added during the last hour of incubation period at 37°C and OD^570^ was measured in an ELISA plate reader. A plot of OD^570^ against cell concentration was constructed and linear regression performed. The equation of the linear plot [Cell concentration (y) = 304.3 x OD^570^ - 330.5] was used to calculate cell numbers of FLS in treated groups. Percentage cell cytotoxicity was calculated by subtracting the cell concentration (y) in a treated well from the mean cell concentration (x) of the untreated respective FLS type cultured in triplicates, and dividing the result by the mean basal FLS cell concentration (x).

Compared to FLS cultured in medium only (5×10^4^ cells / well in a 96-well plate) TLR2 activation with Pam_3_CSK_4_ 100μg/ml for 24-hours led to increase in cell concentration of both NHS (31.3±5.4 %) and RA FLS (24.1±15.1 %). Figure 1 illustrates that pre-treatment with SMV for 4-hours followed by TLR2 activation, led to cell cytotoxicity of both NHS and RA FLS with modestly significant dose dependence (ANOVA F-statistic = 4.66, *P* = 0.0432). SMV 50μM caused microscopically visible cell death and was therefore considered toxic dose for FLS.

**Figure 1.**
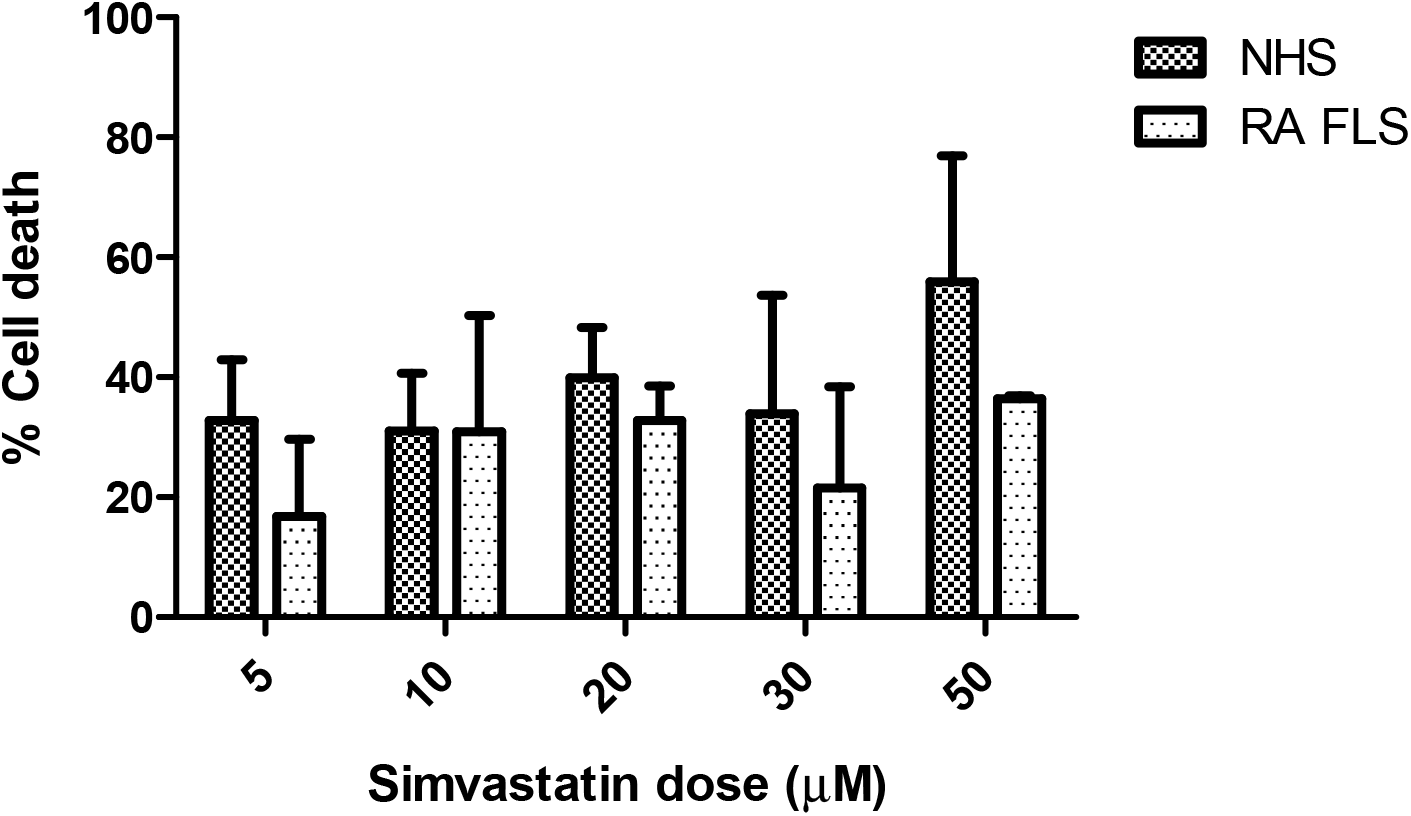
Cell cytotoxicity (%) of FLS. Note: FLS (NHS and RA FLS) were pre-treated with SMV for 4-hours followed by activation with TLR2 ligand, Pam_3_CSK_4_ (100ug/ml), for 24-hours. Cell cytotoxicity was compared to basal levels in medium as measured by Alamar-blue cell cytotoxicity assay.

As Alamar-blue is metabolized in the mitochondria, kinetic recordings of change in OD^570^ can be used as indicator of cellular metabolism. To determine the rate of FLS metabolism, Alamar-blue assay was performed in a kinetic manner at 37°C and absorbance 570nm read every 14 seconds for 1.5-hours in a kinetic ELISA plate reader. First 30-minutes of readings were discarded to allow for cell acclimatization. OD^570^ was plotted against time for each well and a line-of-best-fit calculated. The gradient of this line determined the metabolic rate of FLS (5×10^4^ cells /well) in each well. The mean of the gradients of FLS, cultured in triplicates, was plotted against SMV dose (Figure 2). The dose response for SMV pre-treatment for 4-hours followed by TLR2 activation for 24-hours was an exponential decay curve, demonstrating progressive decrease in metabolic rate with increasing doses of SMV. This data confirmed cell cytotoxicity of SMV in a dynamic manner. It also showed that RA FLS had higher basal metabolic rate than NHS and that TLR2 activation did not significantly alter FLS metabolism (Figure 2).

**Figure 2.**
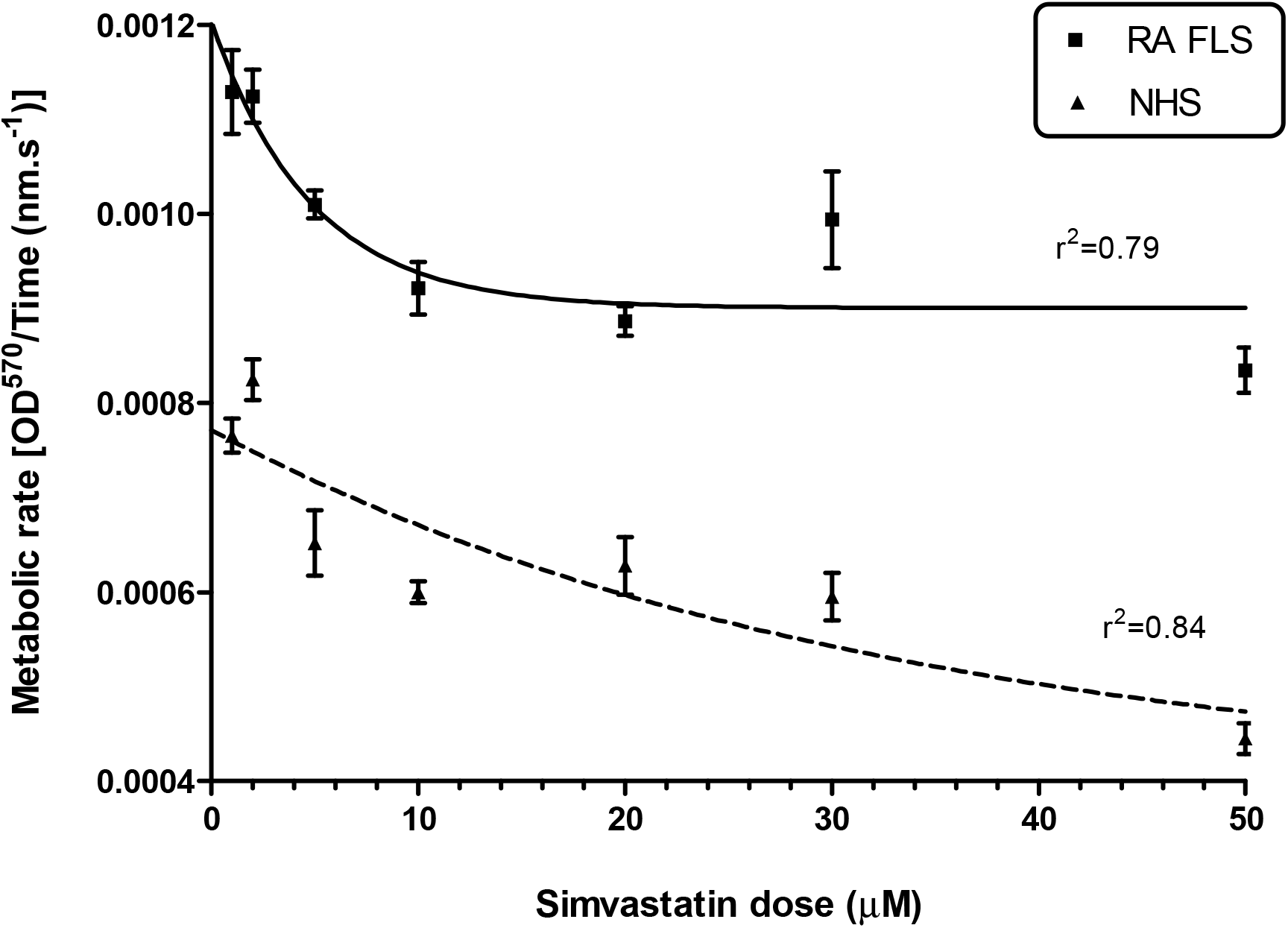
Metabolic rate of FLS determined by Alamar-blue kinetic assay. Note: Dose 1 represents untreated (medium) FLS and dose 2 represents Pam_3_CSK_4_ only (100μg/ml) treated cells.

### Intracellular ROS production

Major sites of cellular reactive oxygen species (ROS) generation include the mitochondrial electron transport chain, the cytosolic NADPH oxidase (NOX) complex and the endoplasmic reticulum. Superoxide is the main initial ROS, which also drives the Fenton reaction derived ROS production (OH^-^ and H_2_O_2_) [3]. NHS and RA FLS treated, in triplicate, with hydrogen peroxide (H_2_O_2_ 20μM) served as positive controls for their respective cell lines. Treatment with H_2_O_2_ for 10 minutes caused 86.4% NHS to produce superoxide radical and 83.5% RA FLS. SMV pre-treatment and TLR2 ligand activation caused superoxide production ranging from 4% to 14% (Figure 3 A-D). Increasing doses of SMV caused greater percentage of TLR2 activated FLS to produce superoxide.

**Figure 3.**
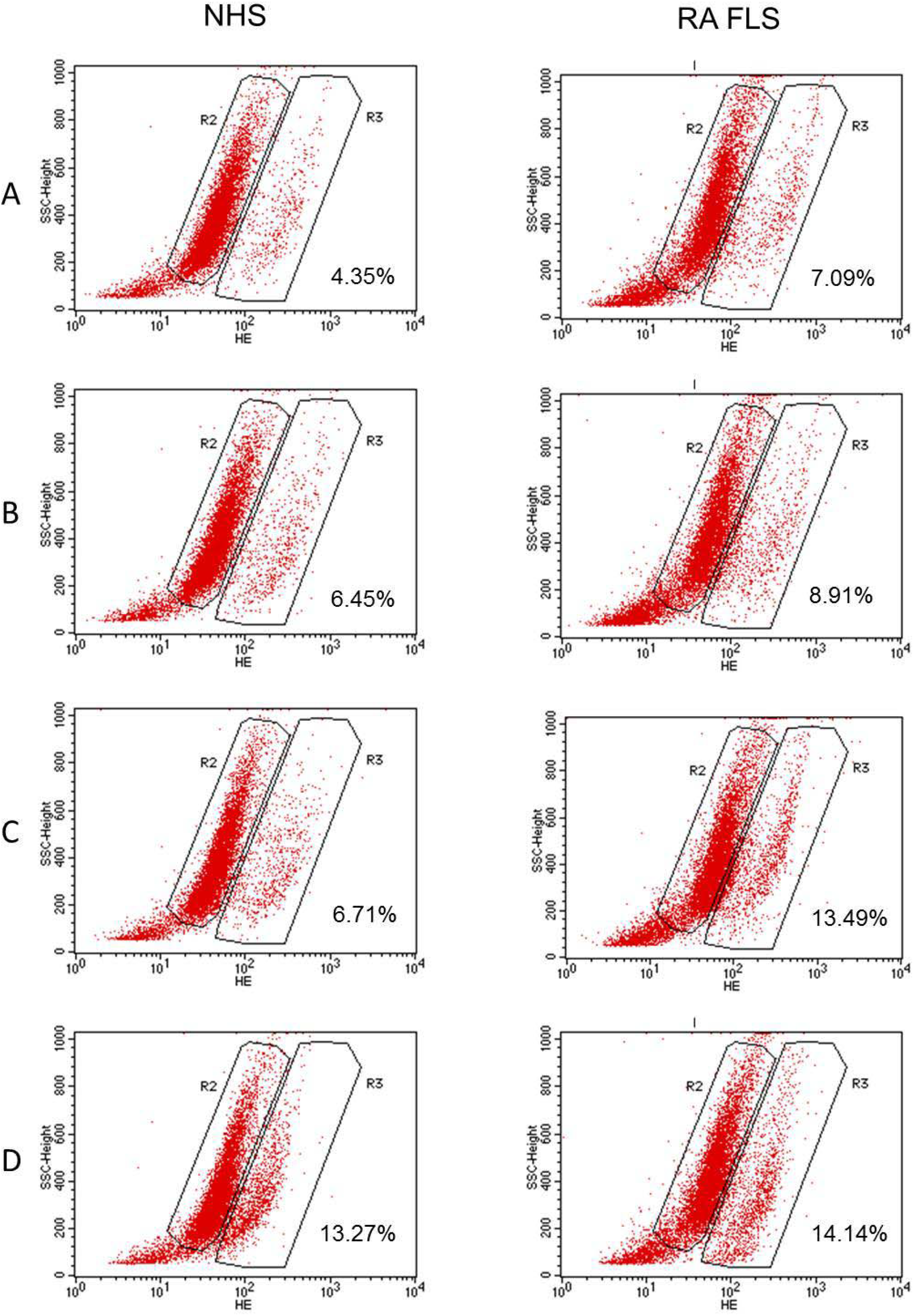
ROS production by FLS as measured by dihyroethidine flow cytometry.

### ROS associated gene expression

To investigate the mechanism of ROS production we carried out a focused gene expression array analysis. NHS and RA FLS were treated in triplicates as four separate groups: (a) synoviocyte medium alone, (b) SMV 30μM for 4-hours, (c) Pam_3_CSK_4_ 100μg/ml for 24-hours and (d) pre-treatment with SMV 30μM for 4-hours followed by Pam_3_CSK_4_ 100μg/ml activation for 24-hours. RNA was extracted from each well and triplicates were pooled in the respective groups. Real-time PCR was performed to assess the fold change in 84-human “oxidative stress and antioxidant defense” genes (SA Biosciences PAHS-065).

Expression of thirteen genes was found to be significantly altered due to one or more treatments compared to medium only basal conditions (Table 1). Three interesting gene expression profiles were noted (Table 1). BCL2/adenovirus E1B 19kDa interacting protein-3 (*BNIP3*), glutathione reductase (*GSR*), neutrophil cytosolic factor-1 (*NCF1*), peroxiredoxin-1 (*PRDX1*), thioredoxin reductase-1 (*TXNRD1*) were significantly upregulated in RA FLS in response to SMV alone, Pam_3_CSK_4_ or their combination as in group-d. Expression of these genes remained essentially unchanged in NHS, while at baseline it was significantly higher than in RA FLS (group a). Therefore SMV with or without TLR2 activation increased the expression of genes in set-1 (Table 1) to the level of NHS. One exception was *NCF1*, which is a component of NADPH oxidase. It was 5-fold lower in RA FLS than NHS at baseline, however its expression was upregulated over 50-fold on pre-treatment with SMV followed by TLR2 activation.

Genes in set-2 had equal expression in NHS and RA FLS at baseline and were significantly upregulated in both cell lines by group-d treatment (SMV+TLR2 activation) however, SMV or Pam_3_CSK_4_ alone did not suffice. These genes included metallothionein-3 (*MT3*), prostaglandin-endoperoxide synthase-2 (*PTGS2* or *COX2*) and superoxide dismutase-2 (*SOD2*). *PTGS2* was unique in that it was upregulated by SMV alone as well, but not by Pam_3_CSK_4_.

**Table 1.**
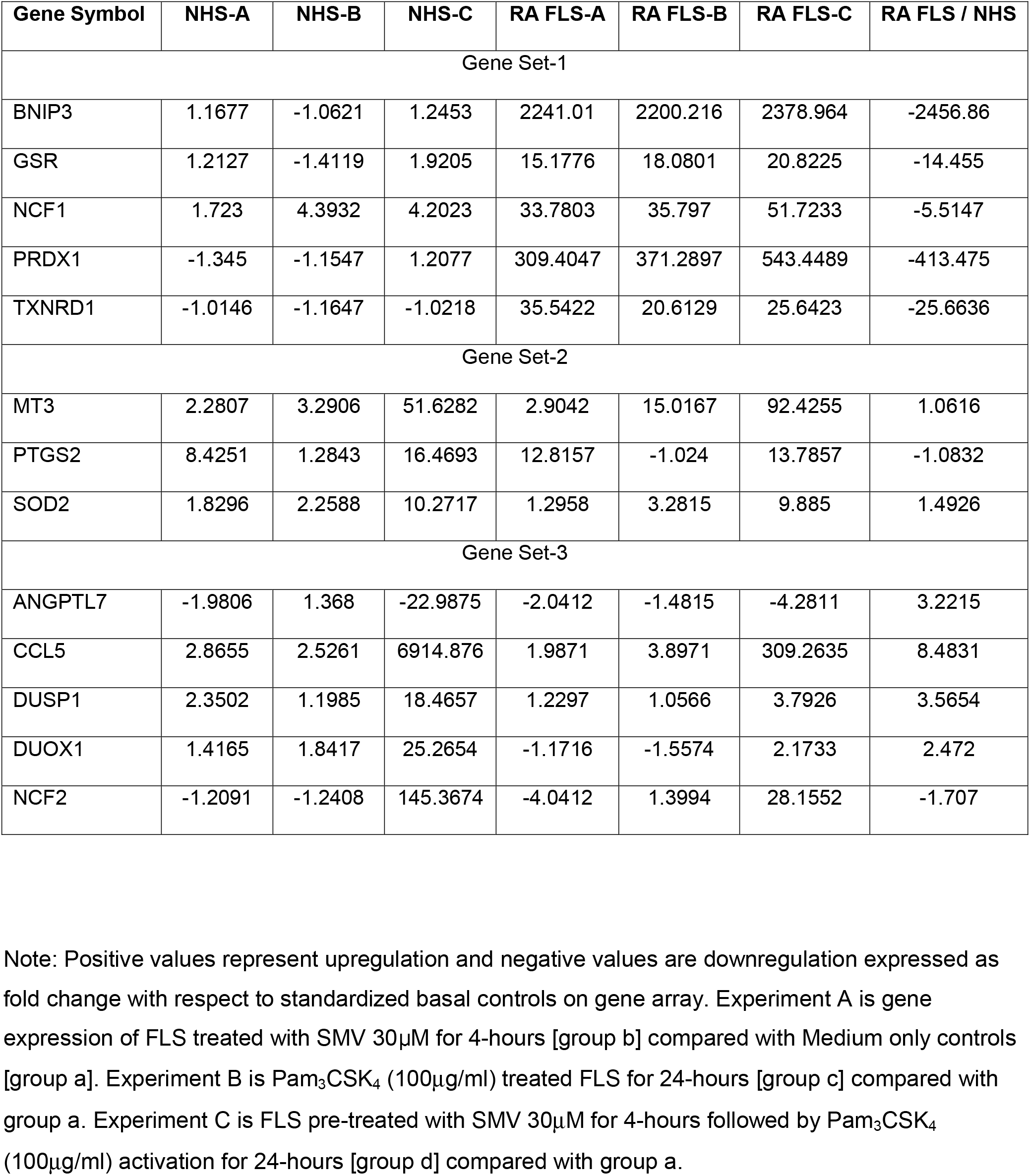
Fold changes of significantly altered ROS associated genes in FLS.

| Gene Symbol | NHS-A | NHS-B | NHS-C | RA FLS-A | RA FLS-B | RA FLS-C | RA FLS / NHS |
| --- | --- | --- | --- | --- | --- | --- | --- |
| Gene Set-1 |  |  |  |  |  |  |  |
| BNIP3 | 1.1677 | -1.0621 | 1.2453 | 2241.01 | 2200.216 | 2378.964 | -2456.86 |
| GSR | 1.2127 | -1.4119 | 1.9205 | 15.1776 | 18.0801 | 20.8225 | -14.455 |
| NCF1 | 1.723 | 4.3932 | 4.2023 | 33.7803 | 35.797 | 51.7233 | -5.5147 |
| PRDX1 | -1.345 | -1.1547 | 1.2077 | 309.4047 | 371.2897 | 543.4489 | -413.475 |
| TXNRD1 | -1.0146 | -1.1647 | -1.0218 | 35.5422 | 20.6129 | 25.6423 | -25.6636 |
| Gene Set-2 |  |  |  |  |  |  |  |
| MT3 | 2.2807 | 3.2906 | 51.6282 | 2.9042 | 15.0167 | 92.4255 | 1.0616 |
| PTGS2 | 8.4251 | 1.2843 | 16.4693 | 12.8157 | -1.024 | 13.7857 | -1.0832 |
| SOD2 | 1.8296 | 2.2588 | 10.2717 | 1.2958 | 3.2815 | 9.885 | 1.4926 |
| Gene Set-3 |  |  |  |  |  |  |  |
| ANGPTL7 | -1.9806 | 1.368 | -22.9875 | -2.0412 | -1.4815 | -4.2811 | 3.2215 |
| CCL5 | 2.8655 | 2.5261 | 6914.876 | 1.9871 | 3.8971 | 309.2635 | 8.4831 |
| DUSP1 | 2.3502 | 1.1985 | 18.4657 | 1.2297 | 1.0566 | 3.7926 | 3.5654 |
| DUOX1 | 1.4165 | 1.8417 | 25.2654 | -1.1716 | -1.5574 | 2.1733 | 2.472 |
| NCF2 | -1.2091 | -1.2408 | 145.3674 | -4.0412 | 1.3994 | 28.1552 | -1.707 |

Note: Positive values represent upregulation and negative values are downregulation expressed as fold change with respect to standardized basal controls on gene array. Experiment A is gene expression of FLS treated with SMV 30 μM for 4-hours [group b] compared with Medium only controls [group a]. Experiment B is Pam<sub>3</sub>CSK<sub>4</sub> (100 μg/ml) treated FLS for 24-hours [group c] compared with group a. Experiment C is FLS pre-treated with SMV 30 μM for 4-hours followed by Pam<sub>3</sub>CSK<sub>4</sub> (100 μg/ml) activation for 24-hours [group d] compared with group a.

Set-3 genes were slightly upregulated in RA FLS at baseline and were only modestly over-expressed on group-d treatment. However, their over-expression was vigorous in NHS on pre-treatment with SMV followed by TLR2 activation. SMV or Pam_3_CSK_4_ alone did not result is substantial upregulation. These genes included chemokine (C-C motif) ligand-5 (*CCL5* or *RANTES*), dual specificity phosphatase-1 (*DUSP1*), dual oxidase-1 (*DUOX1*) and neutrophil cytosolic factor-2 (*NCF2*). Both *DUOX1* and *NCF2* are components of NADPH oxidase. Angiopoietin-like-7 (*ANGPTL7*) is also included in this group because its pattern was similar but in opposite direction i.e. it was downregulated.

### Apoptosis

The 2200-fold upregulation of *BNIP3*, a pro-apoptotic protein, in RA FLS led us to investigate apoptosis as a mechanism of observed cell cytotoxicity in TLR2 activated FLS. SMV as a single treatment has previously been shown to cause apoptosis in NHS [32]. Annexin V flow cytometry assay indeed showed a dose dependent apoptosis with SMV (Figure 4 A-D). It also confirmed that RA FLS had lower basal apoptosis (4.3%) than NHS (14%) [33]. SMV pre-treatment for 4-hours ameliorated this difference and increased apoptosis in both cell lines. The extent of apoptosis was sufficient to explain the cell cytotoxicity observed with the Alamar-blue assay.

**Figure 4.**
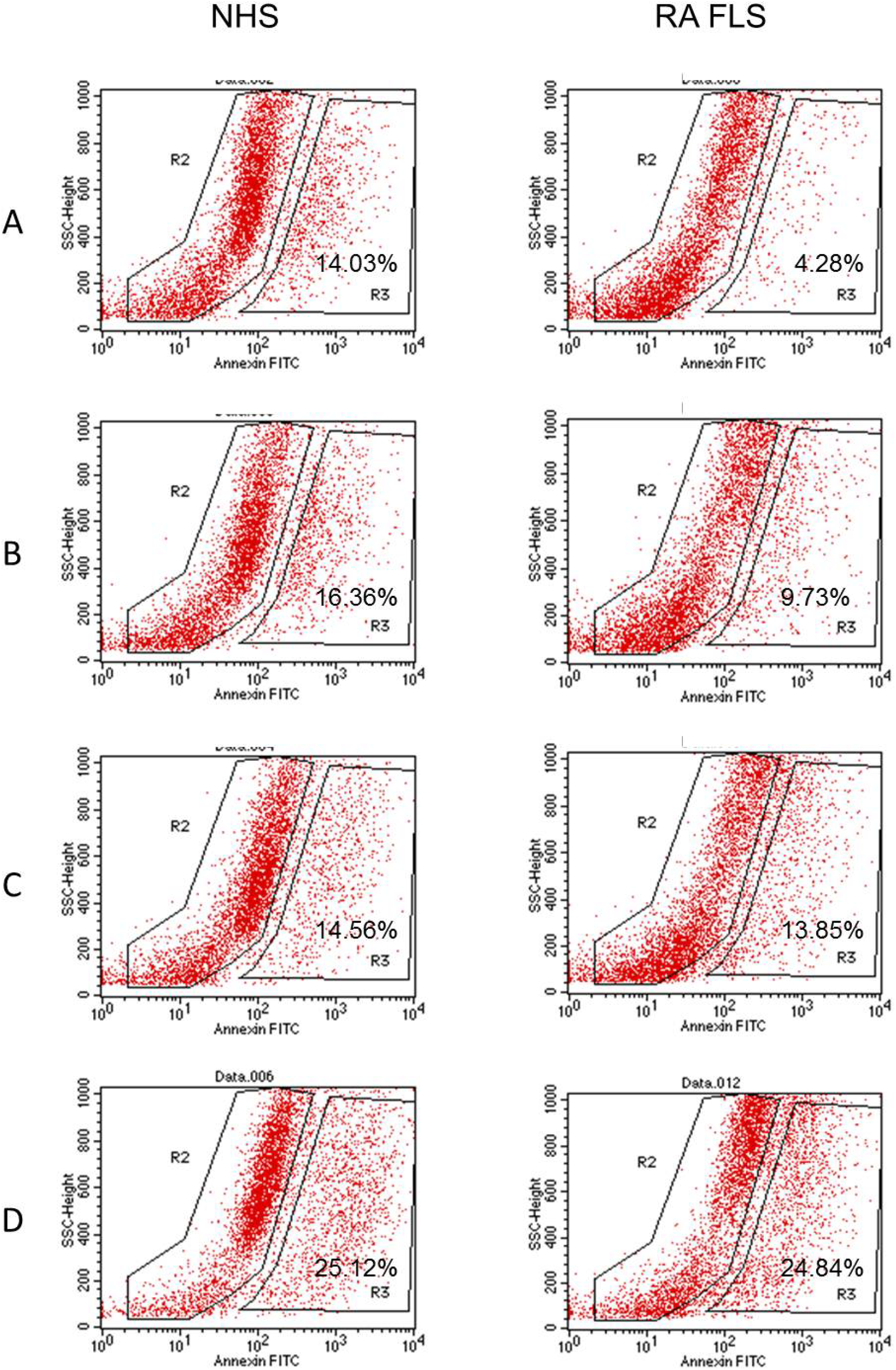
Apoptosis of FLS as measured by Annexin V flow cytometry.

### Cytokine ELISA

Pro-inflammatory cytokine TNF-α is known to cause apoptosis [2]. ELISA for TNF-α showed absence of any TNF-α production by both cell lines in basal and treated conditions i.e. with TLR2 ligand and SMV pre-treatment (data not shown). However, IL-6 was produced by both NHS and RA FLS. As shown in Figure 5, RA FLS produced more IL-6 (171pM) than NHS (128pM) at baseline (*P*=0.09) and responded to TLR2 ligand treatment with significantly higher IL-6 production (T-test=4.72, *P*=0.0092). Interestingly, SMV pre-treatment caused a dose dependent increase in IL-6 production in both cell lines compared to basal IL-6 production (F = 57.29, *P*<0.0001). Both NHS and RA FLS produced statistically similar levels of IL-6 in response to SMV pre-treatment and TLR2 activation (P<0.05).

**Figure 5.**
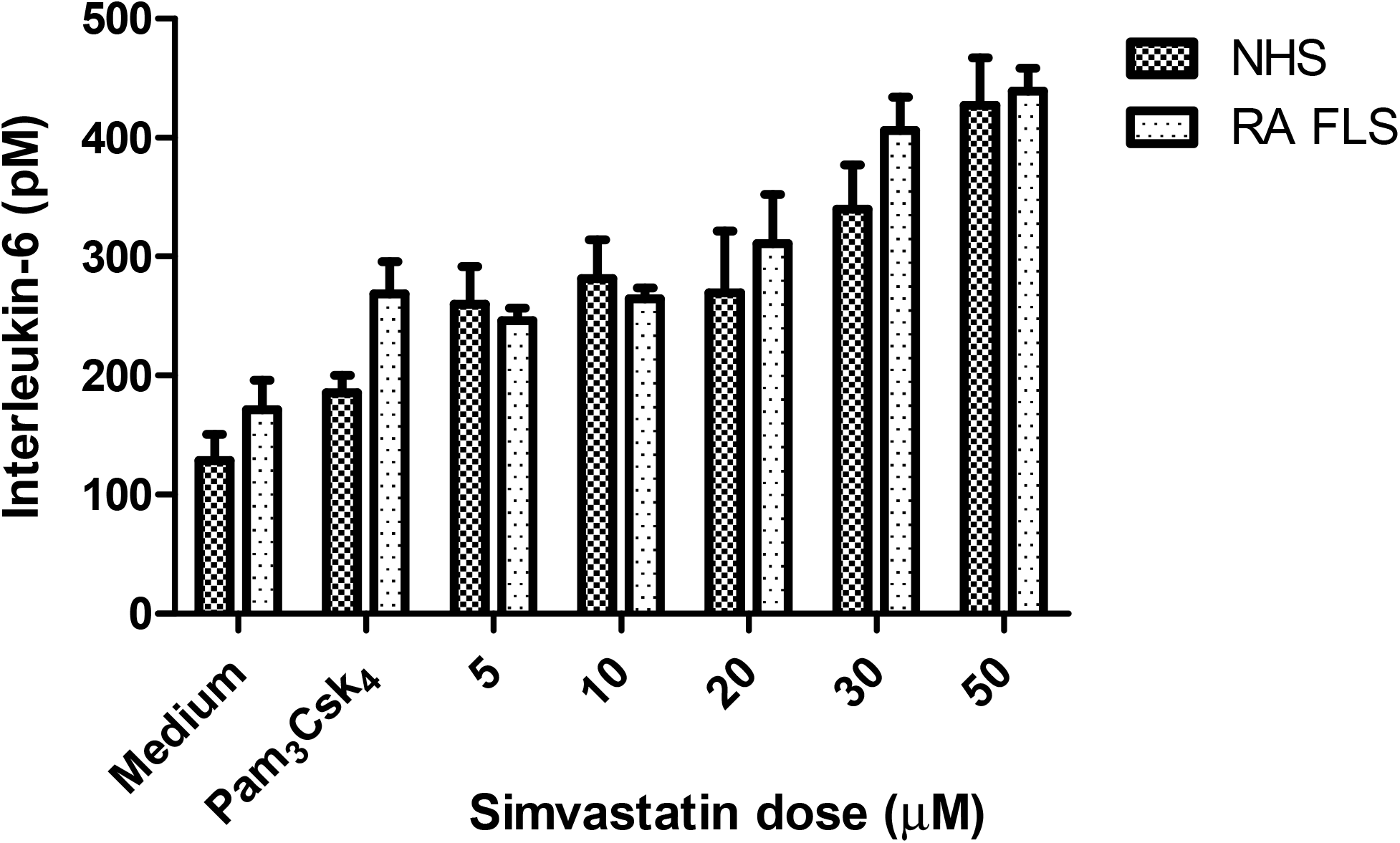
Interleukin-6 production by FLS in response to SMV pre-treatment followed by TLR2 activation.

### Akt and BNIP3 Western blots

Increasing levels of IL-6 are indicative of nuclear factor kappa-B (NF-кB) activation which provides a cell survival signal. The serine–threonine kinase Akt can promote NF-кB activity [34]. NHS and RA FLS were treated in triplicates with a) medium only, b) Pam_3_CSK_4_ 100μg/ml, c) pre-treated with SMV 10μM for 4-hours followed by stimulation with Pam_3_CSK_4_ (100μg/ml) for 24-hours and d) pre-treatment with SMV 30μM and TLR2 activation as in group c. The Western blot for Akt in extracted protein from pooled triplicates showed a differential expression in RA FLS compared to NHS (Figure 6). Akt levels were lower in RA FLS and decreased further with SMV pre-treatment, becoming absent with 30μM dose. In NHS, Akt levels showed an increasing trend with SMV pre-treatment (Figure 6). Thus, in RA FLS, Akt is likely to play a significant role in mediating SMV induced apoptosis.

**Figure 6.**
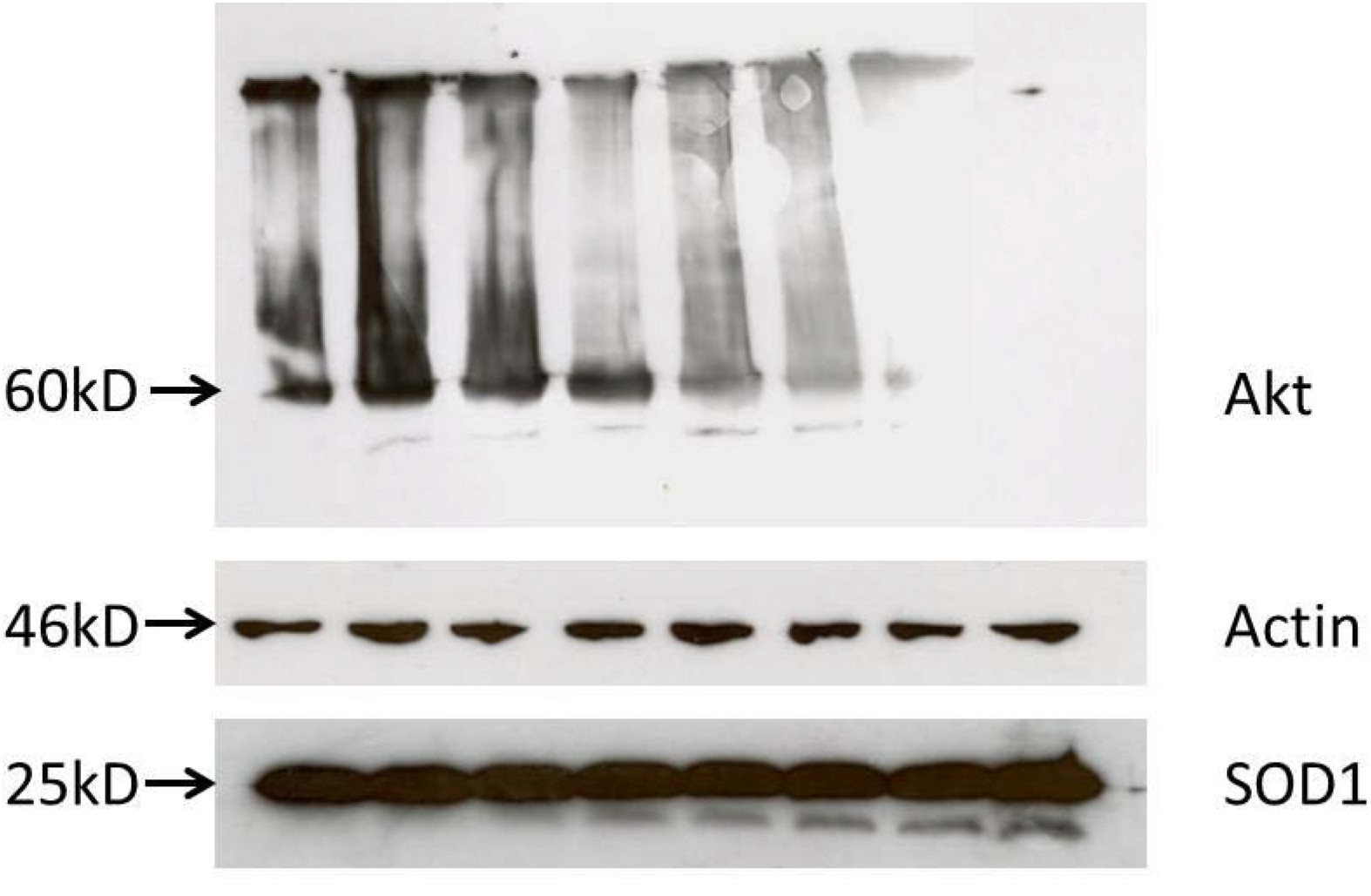
Western blots of Akt, SOD1. Note: First four bands are for NHS, followed by four bands for RA FLS in order: (i) Medium control, (ii) Pam_3_CSK_4_ (100μg/ml), (iii) Pre-treatment with SMV 10μM followed by TLR2 activation, (iv) Pre-treatment with SMV 30μM followed by TLR2 activation. BNIP3 was not detected.

We also performed western blot for BNIP3 as it was found to be highly upregulated in RA FLS. However, no BNIP3 protein was detected by western blot, likely due to low BNIP3 concentration and sensitivity of the technique. SOD1 was easily detectable and its levels were found to remain unchanged irrespective of cell line and treatment, in agreement with gene expression data. Actin was used as background control.

### Inositol phosphate (IP) levels

IPs compete with phosphotidylinositols (PIs e.g.PIP_3_) to inhibit the Akt pathway, leading to apoptosis [30]. Thus, we investigated the effect of TLR2 activation and SMV pre-treatment on IP (1-4) levels in FLS. It has been previously shown in multiple cell lines that in apoptosis IP1 and IP2 levels increase whereas, IP3 and IP4 levels decrease [29, 30]. We found that IP1 (F=6.61, *P*=0.0041), IP2 (F=4.36, *P*=0.02) and IP3 (F=10.54, *P*=0.0005) levels were significantly decreased after pre-treatment with SMV 30μM and TLR2 activation in both NHS and RA FLS (Figure 7 A-C). However, IP4 levels (F=0.82, *P*=0.5) remained unchanged (Figure 7D). Both cell lines produced statistically similar levels of IPs (1-4) at baseline and with each treatment (P<0.05).

**Figure 7.**
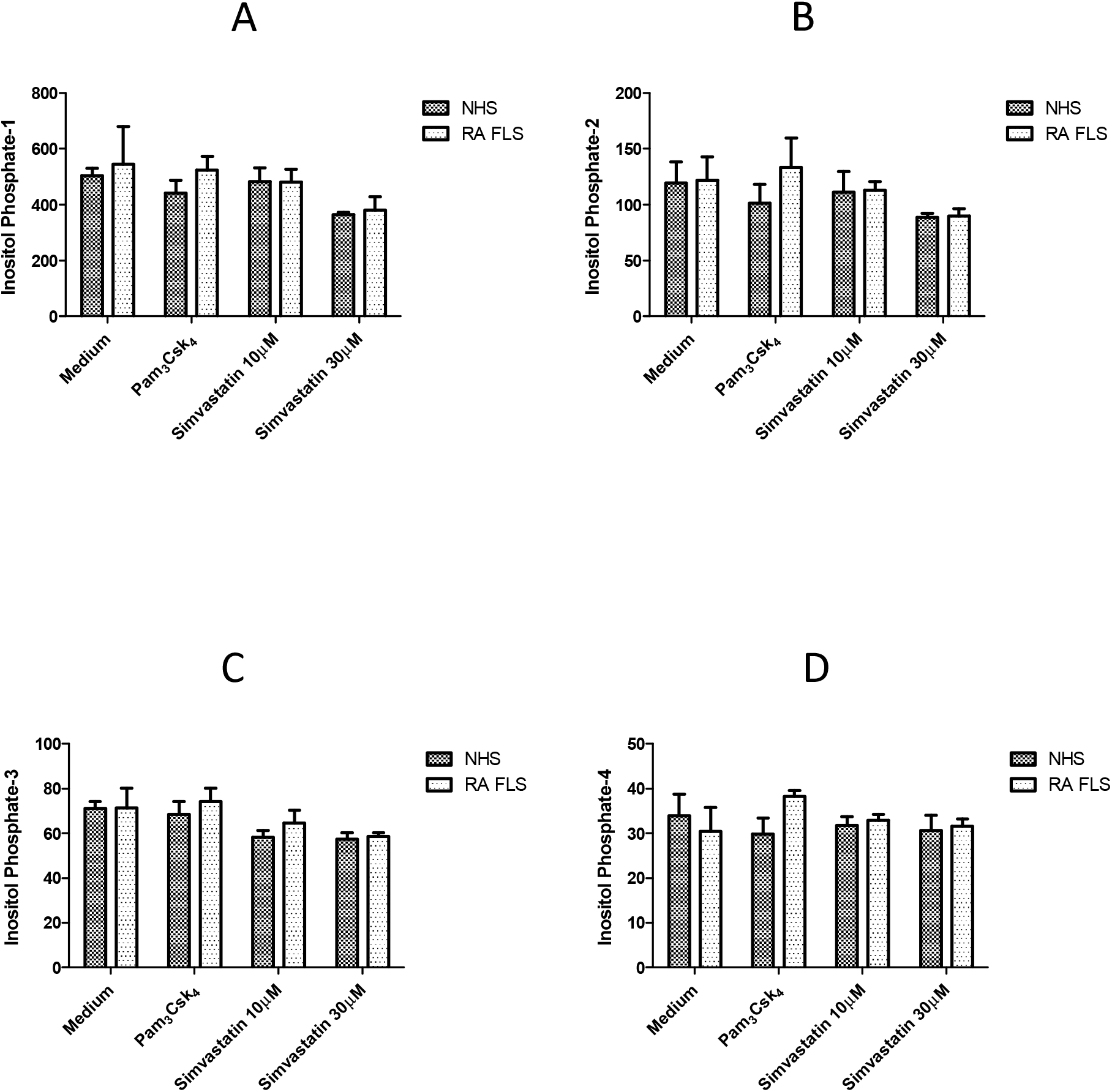
Ionositol phosphate levels in FLS pre-treated with SMV followed by TLR2 activation.

## Discussion

In RA the intimal lining layer expands due to proliferation of FLS, which are major effectors of cartilage destruction in RA [1-4] and the focus of this study. A key role for TLR2 activation of FLS in RA pathogenesis is supported by multiple lines of evidence. TLR2 activation in FLS results in induction of various cytokines, angiogenic factors and matrix-metalloproteinases which subsequently lead to cartilage damage [7, 35]. TLR2 is expressed at high levels in the synovial intimal lining comprised of FLS and also at sites of pannus invasion into cartilage and bone [2, 7]. TLR2 knockout mice are protected from joint inflammation [8], while intra-articular injection of the TLR2 led to development of destructive arthritis in mice [8]. It was recently demonstrated in RA synovial explant cultures that TLR2 agonist Pam_3_CSK_4_, upregulated pro-inflammatory cytokine production in RA mononuclear cells, an effect that was significantly blocked by TLR2 monoclonal antibody, OPN301 [36].

SMV was shown to inhibit the incidence and severity of collagen induced arthritis (CIA) in DBA/1 mice when administered prophylactically, following intraperitoneal type II collagen challenge [13]. Similar effect was observed when SMV was given therapeutically after the development of CIA in mice. In our in vitro studies, we preferred to employ the prophylactic experimental model to minimize direct chemical effects of the drug, such as potential extracellular scavenging of ROS, and to more clearly observe its immunomodulatory properties in FLS. After pre-treatment with SMV, FLS were activated with the TLR2 agonist, Pam_3_CSK_4_, to more closely mimic the *in vivo* processes of RA pathogenesis.

We report here that SMV in TLR2 activated FLS causes cell cytotoxicity in normal (NHS) and RA FLS mediated by apoptosis in a dose dependent manner. Mitochondrial metabolism of FLS also decreased dose dependently. Both mitochondrial metabolism and apoptosis in RA FLS was lower (4.3%) that NHS (14%) at baseline pointing to the more aggressive phenotype of RA FLS. Superoxide production was also enhanced in both cell lines and this may contribute to FLS apoptosis. SMV modulates several ROS associated genes including NADPH oxidase components such as DUOX1, NCF1 and NCF2, which may augment superoxide production.

There was differential expression on BNIP3 in NHS and RA FLS, in that it was upregulated on treatment (SMV, TLR2 activation and both) in RA FLS only (Table 1). Overexpression of BNIP3 is known to induce FLS apoptosis that is inhibited by proinflammatory cytokines [37]. BNIP3 was found to be widely expressed in RA synovium [37]. We found that SMV alone or in combination with TLR2 ligand upregulates BNIP3 mRNA in RA FLS > 2200-folds. However, we were unable to detect BNIP3 on western blot analysis likely due to low titers of the protein. BNIP3 interacts with BCL2, an anti-apoptotic protein and inhibits it. Interestingly BNIP3 is known to induce apoptosis, even in the presence of BCL2 [38]. Thus BNIP3 can potentially cause FLS apoptosis independent of the activation of Akt and NF-кB. Moreover, Akt levels were decreased in RA FLS which could also contribute to SMV mediated dose dependent apoptosis in this cell line (Figure 6).

Besides cell death signals we simultaneously identified cell survival signals in both cell lines on pre-treatment with SMV and TLR2 activation. Interestingly, SMV had similar effects on normal and RA FLS except for Akt levels and expression of set-1 genes (Table 1). Upregulation of anti-oxidant enzymes and proteins in RA FLS such as GSR, MT3, PRDX1, SOD2 and TXNRD1 may serve to limit ROS associated cellular damage. RANTES (CCL5), which is a potent chemokine for monocytes and memory T helper cells [2], was upregulated ∼ 300 fold in RA FLS and over 6000-fold in NHS after SMV pre-treatment and TLR2 activation. RANTES overexpression would lead to potentiation of the inflammatory response.

IL-6 levels were markedly increased (Figure 5). The maximum levels of IL-6 were detected in supernates from NHS and RAFLS treated with a toxic dose (50μM) of SMV. It has previously been shown that 24-hour incubation with SMV 50μM in the presence of IL-1β caused inhibition of NF-кB [39]. As shown here by cell cytotoxicity and FLS mitochondrial metabolism data, pre-treatment with SMV 50μM for 4-hours followed by complete washing and TLR2 activation of FLS caused ∼ 60% cell death in NHS and ∼ 40% in RA FLS (Figure 1). Based on this data and light microscopy we considered this dose of SMV to be toxic to FLS.

SMV was shown to reduce serum IL-6 levels in DBA/1 mice [13]. In an animal model of CIA IL-6 production can be from many different cell types including macrophages. In the TARA study the patients were treated with atorvastatin in addition to other DMARDs including methotrexate which has been shown to lower IL-6 levels [20, 40]. Thus, the effect of statin therapy on IL-6 may be confounded by use of other DMARDs in that study. In our in vitro model of FLS, SMV pre-treatment followed by TLR2 activation caused a dose dependent increase in IL-6, a cytokine that is produced as a result of NF-кB activation [2]. This indicates that SMV may cause apoptosis through NF-кB independent pathway(s) and FLS react to survive by activating NF-кB.

Akt pathway activation promotes NF-кB activity [34] and increased thickness of the Akt band was observed in NHS on SMV pre-treatment, in a dose dependent manner (Figure 6). Moreover, inositol phosphates can competitively block Akt by binding to the PH domain of Akt, leading to apoptosis [29]. Previously, we have shown in multiple cell lines that increased levels of IP1 and IP2 correlate with apoptosis [29, 30]. However, IP1, IP2 and IP3 levels in both FLS types were significantly decreased after pre-treatment with SMV in a dose dependent manner, indicating a cell survival signal. The decrease in the levels of IPs may not have been sufficient to disinhibit Akt in RA FLS.

The results of this study help to reconcile the discrepancies observed in the literature for treatment of RA with statins [13, 17-27]. We show here that SMV in the presence of TLR activation causes apoptosis in RA FLS possibly by increasing ROS production, upregulation of BNIP3 and inhibition of Akt. FLS react to survive by upregulating anti-oxidant genes, increasing IL-6 and decreasing inositol phosphates, which are competitive inhibitors of Akt. These effects in addition to upregulation of RANTES may worsen RA disease activity and may have led to the absence of beneficial effect in other large cohorts [25, 26].

## Conclusions

We carried out an in vitro study to investigate the immunomodulatory effects of SMV in TLR2 activated of normal and RA FLS. We found that SMV pre-treatment, in general, had similar effects on both cell types. We observed dose dependent cell cytotoxicity of FLS mediated by apoptosis. This may be mediated by increased ROS production, upregulation of BNIP3 and inhibition of Akt in FLS. At the same time we observed cell survival signals such as upregulation of anti-oxidant genes, increased levels of IL-6 and decreased inositol phosphates. We hypothesize that SMV induces apoptosis in FLS through NF-кB independent pathway(s) and FLS react to survive by activating NF-кB. Our investigations help to reconcile the discrepancies observed in the literature for treatment of RA with statins. More potent inhibition of cell survival signals by other DMARDs may allow simvastatin to be used as adjunctive therapy in RA.

## Acknowledgment

This work was supported by Arkansas Biosciences Institute, Tobacco Award 2010, ARIA # 37501, UAMS (MS and RAO) and by the funds provided by the Department of Internal Medicine and the Division of Rheumatology, UAMS. Authors wish to thank Dr. Krishnaswamy Kannan for his support with flowcytometry experiments and culture setups and Ms. Preeti Tripathi for technical assistance in RNA extraction and cDNA preparation and the Department of Veteran Affairs (CAVHS-Little Rock) for the permission to use ABI-7900 HT and FACS Calibur machines.

## Abbreviations

RA: rheumatoid arthritis
FLS: fibroblast-like synoviocytes
Simvastatin: SMV
ROS: reactive oxygen species
DMARD: disease modifying anti-rheumatic drug

## Authors’ contributions

MS designed the study, performed majority of the experiments, data analysis, and manuscript preparation. Western blots and Inositol phosphate experiments were performed by ZK and NA. Funding was in part provided by a grant from Arkansas Biosciences Institute awarded to RAO as PI who also provided the research setup for all experiments conducted by MS. All authors read and approved the final manuscript.

